# Environmentally driven changes in leaf spectral signatures impact plant spectral taxonomy

**DOI:** 10.1101/2023.05.09.538942

**Authors:** Natalia L. Quinteros Casaverde, Douglas C. Daly, Shawn P. Serbin

## Abstract

Plant identification is crucial for the conservation and management of natural areas. The shortwave spectral reflectance of leaves is a promising tool for rapidly identifying plants at different taxonomic levels. However, leaf spectral reflectance changes in response to biotic and abiotic conditions. Here we assess whether this variability in spectral reflectance affects the accuracy of classification methods currently used to predict plant taxonomy and identify factors that most influence leaf spectral signatures, as proxies for predicted biochemical and structural traits. We used leaf reflectance from 42 woody species from the living collection at the New York Botanical Garden across two sets of pairwise samplings (spring 2019/summer 2020 and spring 2019/winter 2021). We found that classification accuracy was poor when only spring samples were used to train models but improved when natural variation across all seasons was incorporated into classification models. To evaluate the influence of species relatedness or growth conditions (temperature, relative humidity, and daylength) on spectrally predicted traits, we applied Partial Least Squares Regression (PLSR) coefficients derived from NEON data to predict foliar traits including photosynthetic pigments, water content, leaf dry mass per area, and carbon and nitrogen content. Results showed that trait variation was not influenced by phylogeny but was significantly influenced by environment, except for water-related spectral bands for species remeasured in summer 2020. These results demonstrate that classification methods developed to handle datasets with large collinearities, such as leaf spectral reflectance, underperform when individual spectral variability is caused by environmental factors. This issue must be addressed for future development of remotely sensed taxonomy applications for evergreen broadleaf vegetation.

## 1 Introduction

In an era of accelerating biodiversity loss, the race to identify and catalog Earth’s plant species has never been more urgent—yet traditional taxonomic expertise is vanishing faster than the species themselves. Plant identification underpins biodiversity conservation (Trias-Blasi and Vorontsova, 2015) and ecosystem management (Noss, 2001), making taxonomic knowledge crucial for ecological understanding. As this expertise declines, multiple disciplines—from community ecology to biogeography—face significant challenges (Pyšek et al., 2013). Technological advances have emerged to address this crisis, including DNA barcoding (Dormontt et al., 2018), environmental DNA (Clare et al., 2022), and leaf spectroscopy (Cavender-Bares et al., 2025; Durgante et al., 2013; Meireles et al., 2020).

Leaf spectral signatures—nature’s optical fingerprints—offer the potential for rapid plant identification. These reflectance patterns across visible to shortwave infrared wavelengths (300-2500 nm) arise because plants have evolved to interact with electromagnetic radiation passing through atmospheric windows (Kiang et al., 2007). Following the Beers-Lambert law, leaf components absorb energy proportionally to their concentration and molar absorptivity (Luther and Nikolopulos, 1913), creating distinctive spectral patterns.

Leaf chemical and structural attributes, including photosynthetic pigments, water content, cellulose, lignin (Brewer, 2012), as well as secondary metabolites (Fine et al., 2021), interact with radiation in characteristic ways. However, these traits demonstrate environmental (Díaz et al., 2016; Lima et al., 2022) and ontological (Chavana-Bryant et al., 2017; Wu et al., 2018, 2017) plasticity, affecting spectral signatures (Castro-Esau et al., 2006; Hesketh & Sánchez-Azofeifa, 2012; Wu et al., 2017). The critical question remains: can spectral identification overcome the challenge of temporal variability to deliver reliable taxonomic results?

This study investigates how seasonal and interannual leaf spectral variability impacts tropical woody species identification in the Enid A. Haupt Conservatory (EAHC) at the New York Botanical Garden. We go beyond simple accuracy metrics to understand the evolutionary and environmental origins of misclassification patterns, revealing whether errors reflect phylogenetic/environmental constraints or methodological limitations. We hypothesize that integrating spectral variability across seasons and years into training datasets will dramatically outperform single-season classification models. Additionally, we employ spectrally-derived leaf traits—including photosynthetic pigments, water, and nitrogen, as well as leaf dry mass per area— as biochemical and structural intermediates to mechanistically link environmental conditions (temperature, humidity, day length) with optical responses. This approach not only explains the physiological drivers of spectral variation but also provides a pathway for predicting and correcting temporal classification challenges in future applications.

## 2 Methods

### 2.1 Data Collection

To evaluate temporal variability effects on spectral-based plant classification, we established a multi-seasonal dataset of 42 woody species (one individual per species) from six botanical families (Acanthaceae (five species), Ericaceae (six species), Fabaceae (eight species), Myrtaceae (five species), Rubiaceae (ten species), and Solanaceae (seven species)) housed in the Rainforest Pavilion at the EAHC (Table 1; Figure S 1). Family selection was based on the availability of multiple woody species for comparative analysis.

**Table 1.** List of 42 species for which spectral data were collected at the Enid A. Haupt Conservatory at the New York Botanical Garden corresponding to six botanical families. Species were sampled in spring 2019, and resampled in the summer 2020 or winter 2021, exact dates of resampling are shown in Table A1.1.

| Family | Scientific Name |
| --- | --- |
| ACANTHACEAE | *Aphelandra flava* Nees; *Aphelandra tetragona* (Vahl) Nees; *Louteridium donnell-smithii* S. Watson; *Megaskepasma erythrochlamys* Lindau; *Thunbergia grandiflora* Roxb; *Thunbergia mysorensis* (Wight) T. Anderson |
| ERICACEAE | *Cavendishia lebroniae* Luteyn; *Macleania bullata* Yeo; *Macleania coccoloboides* A.C.Sm.; *Macleania pentaptera* Hoerold; *Macleania rupestris* (Kunth) A.C.Sm.; *Psammisia ramiflora* Klotzsch |
| FABACEAE | *Calliandra haematocephala* Hassk.; *Calliandra tergemina* (L.) Benth.; *Hymenaea* sp.; *Inga edulis* Mart.; *Mucuna bennettii* F. Muell.; *Pterocarpus officinalis* Jacq.; *Strongylodon macrobotrys* A. Gray; *Tipuana tipu* (Benth.) Kuntze. |
| MYRTACEAE | *Campomanesia guazumifolia* (Cambess.) O. Berg; *Eugenia uniflora* L.; *Pimenta racemosa* (Mill.) J.W. Moore; *Psidium friedrichsthalianum* (O. Berg) Nied.; *Psidium guajava* L. |
| RUBIACEAE | *Cinchona* sp.; *Coffea arabica* L.; *Deppea splendens* Breedlove & Lorence; *Exostema nitens* Urb.; *Hamelia cuprea* Griseb.; *Osa pulchra* (D.R. Simpson) Aiello; *Pogonopus speciosus* (Jacq.) K. Schum.; *Posoqueria latifolia* (Rudge) Roem. & Schult.; *Psychotria nervosa* Sw.; *Rosenbergiodendron longiflorum* (Ruiz & Pav.) Fagerl. |
| SOLANACEAE | *Brunfelsia grandiflora* D. Don; *Brunfelsia nitida* Benth.; *Brunfelsia undulata* Sw.; *Cestrum aurantiacum* Lindl.; *Iochroma cyaneum* (Lindl.) M.L. Green ex G.H.M. Lawr. & J.M. Tucker; *Iochroma fuchsioides* (Bonpl.) Miers; *Solandra* sp. |

Our sampling design captured three distinct seasonal periods across two years. The initial spring collection (days 105-112, 2019) established baseline measurements with five mature and sun-lit leaves per individual for all 42 species. Summer sampling (day 251, 2020) included three leaves (mature and sun-lit) from 25 previously sampled species, while winter collection (day 57, 2021) targeted three leaves (mature and sun-lit) from 17 different species from the original cohort (Figure S 1). During each collection, ontological stage of leaves was visually assessed based on the specific greenness and rigidity characteristics of each species. The reduced sampling intensity in 2020-2021 resulted from COVID-19 restrictions limiting personnel access to the enclosed conservatory, and the necessary steps were taken to ensure appropriate handling of the statistical analysis. All collections occurred between 10:00-15:00 hours to minimize diurnal variation effects.

Spectral measurements employed two calibrated spectroradiometers: a Spectra Vista Corporation (SVC HR 1024i) and Spectral Evolution (PSR+) in 2019, with SVC HR 1024i used exclusively for 2020-2021 collections. Both instruments utilized identical SVC LC-RP-Pro leaf clips with internal, full-spectrum light sources and were calibrated against Spectralon® white references. Since both spectroradiometers covered identical spectral ranges with radiometric calibration and consistent reference standards, no inter-instrument calibration was required (Serbin et al., 2019).

Each leaf underwent 4-6 spectral measurements across the lamina, avoiding major vascular bundles. We utilized the R-FieldSpectra package for data processing, applying splice corrections for FieldSpec4 data to align VIS and SWIR sensors with NIR sensors (Meacham-Hensold et al., 2019), while SVC discontinuities were corrected using vendor software. Quality control procedures included bias thresholds at 450 nm to ensure proper leaf clip contact and removal of replicates deviating more than two standard deviations from the mean. Final spectra represented averaged replicates per leaf, with analysis restricted to 400-2400 nm to avoid low signal-to-noise regions at spectral range edges.

### 2.2 Classification of leaf spectra at different taxonomic levels considering temporal variation

To investigate how temporal changes in leaf spectral signatures impact taxonomic classification performance, we employed two complementary analytical approaches that directly test the effect of temporal variability on model reliability across family, genus, and species levels.

Our "Split" approach simulated real-world temporal generalization challenges by training models exclusively on spring 2019 data and testing on temporally distinct summer 2020 and winter 2021 collections. In contrast, our "Stratified" approach integrated temporal variability into both training and testing phases using stratified random partitioning (70% training/30% testing) across all seasons and years, with sampling balanced by taxonomic level and collection date. Family and species analyses utilized all 42 individuals, while genus-level classification focused on eight genera with multiple congeneric representatives: *Aphelandra, Brunfelsia, Calliandra, Cinchona, Iochroma, Macleania, Psidium*, and *Thunbergia* (Table 1).

We employed Partial Least Squares Discriminant Analysis (PLS-DA) as classification method. PLS-DA is specifically designed for highly collinear data (Lee et al., 2018) such as spectral data, as well as imbalanced datasets commonly found in ecological studies (Lindström et al., 2011; Ruiz-Perez et al., 2020). PLS-DA extends PLS regression for classification tasks by reducing spectral reflectance bands (predictor variables) into latent components that maximize covariance between taxonomic classes and spectral signatures (Djuris et al., 2013). This dimensionality reduction creates linear combinations of original variables optimized for discriminating taxonomic identity rather than simply capturing spectral variance.

The PLS-DA Model optimization employed bootstrap resampling, with 1000 resampling iterations to avoid overfitting, within the train function from the caret package (Kuhn, 2008). We tested up to 30 components with 500 iterations to ensure convergence. The kernel PLS classification method was selected for its nonlinear multivariate robustness with small sample sizes (Allegrini and Olivieri, 2023), and data preprocessing was set to center and scale the data. Performance evaluation included Cohen’s kappa (κ) coefficient for classification accuracy and comprehensive precision, recall, and F1 metrics calculated at both macro (unweighted means penalizing poor minority class performance) and weighted (accounting for class imbalance) levels (Table S 1; Mathew et al., 2023).

### 2.3 Detection of variability within spectral reflectance

To identify wavelength regions exhibiting high temporal variability within leaf spectral signatures, we calculated coefficients of variation (CV) between spring 2019 baseline measurements and summer 2020 or winter 2021 remeasurements. CV calculations were performed separately for each resampling season due to different species compositions in the summer and winter datasets. We then calculated overall mean CV per waveband to identify spectral regions with consistently high temporal variation across species. Spectral signatures and resulting CV patterns were visualized using heatmaps generated with the Heatmap function from the ComplexHeatmap package (Gu, 2022), enabling identification of wavelength-specific temporal variation patterns across species and seasons.

### 2.4 Prediction of Leaf Structural and Chemical Traits

To test whether leaf spectral reflectance was affected by environmental factors, we predicted six traits from leaf optical properties: (1) total chlorophylls A&B (𝜇g/mL), (2) chlorophyll A (𝜇g/mL), (3) carotenoids (𝜇g/mL), (4) equivalent water thickness (EWT, g/m²), (5) nitrogen content (%), and (6) leaf dry mass per area (LMA, g/m²). We employed a Partial Least Squares (PLS) regression framework implemented by Burnett et al. (2021) using the R packages spectratrait (Serbin et al., 2025) and pls (Mevik & Wehrens, 2007).This framework predicts leaf traits using regression coefficients by waveband, which quantify the contribution of each spectral band to the prediction of each specific trait (Burnett et al., 2021; Serbin et al., 2014).

To obtain PLS regression coefficients, we utilized the National Ecological Observatory Network’s field spectral and foliar trait databases (National Ecological Observatory Network (NEON), 2025a, 2025b) from six sites representing Arctic, temperate, and tropical ecosystems: HEAL (Healy, Alaska), GUAN (Guánica Forest, Puerto Rico), WREF (Wind River Experimental Forest, Washington), SERC (Smithsonian Environmental Research Center, Maryland), RMNP (Rocky Mountain National Park, Colorado), and PUUM (Pu’u Maka’ala Natural Area Reserve, Hawaii). For the step-by-set procedure see Quinteros Casaverde and Serbin (2025). We applied these coefficients to the NYBG dataset and constructed jackknife confidence intervals to assess prediction precision (Burnett et al., 2021; Kathuria et al., 2025; Serbin et al., 2019, 2014).

### 2.5 Effects of the EAH Conservatory environment over the predicted traits

#### 2.5.1 Environmental variables

To assess environmental effects on spectrally predicted traits (photosynthetic pigments, EWT, LMA, and percent of nitrogen) we obtained environmental data from sensors within the Conservatory connected to a PRIVA® system that recorded temperature and relative humidity every five minutes. We focused on these two variables because of their fundamental importance for plant physiology (Driesen et al., 2020; Lysenko et al., 2023) and their combined influence on vapor pressure deficit, which directly affects plant water relations and gas exchange (Anderson, 1936). Wind speed and water availability were treated as constant parameters with unlimited availability and consequently were excluded from the model, as these conditions remained stable throughout the study period.

Day length calculations were performed using the daylength function from the insol package (Corripio, 2019), while environmental summaries included the mean, standard deviation, and range of temperature and relative humidity for the 30-day period prior to and including each measurement day. This 30-day window was selected to capture recent environmental conditions that could influence plant physiological responses (Van Goethem et al., 2013).

#### 2.5.2 Predicted trait variability assessment

To quantify shifts in predicted traits between baseline measurements and subsequent remeasurements (2020-2021), we employed appropriate paired comparisons based on data distributions. For normally distributed traits, we conducted paired t-tests using the t.test function, while non-parametric traits were analyzed using Wilcoxon signed-rank tests via the wilcox.test function, both from the R stats package (R Core Team, 2013). Effect sizes were calculated as Cohen’s d (mean difference divided by standard deviation) to standardize the magnitude of temporal shifts across different trait scales.

To visualize multivariate separation between baseline and remeasurement trait assemblages, we performed linear discriminant analysis (LDA) using the lda function from the MASS package (Venables and Ripley, 2002). The resulting first two discriminant axes were plotted to represent the multidimensional trait space, enabling assessment of overall trait compositional shifts between temporal sampling points while accounting for trait covariance structure.

#### 2.5.3 Predicted trait phylogenetic signal analyses (PSA)

To determine whether taxonomic family should be included as a random factor in our mixed-effects models, we quantified the contribution of plant evolutionary history to spectral signature variability. This analysis allowed us to understand whether closely related species exhibited similar spectral traits due to shared evolutionary history, which would necessitate accounting for phylogenetic non-independence in our models.

Before the PSA, we first addressed temporal variability by partitioning species based on resampling dates (e.g., spring 2019 vs. summer 2020, spring 2019 vs. winter 2021) and calculating interquartile ranges for each trait within these temporal groups. This approach allowed us to separate phylogenetic effects from seasonal variation in trait expression while having enough data to detect a phylogenetic effect in a continuous trait (Yao and Yuan, 2025).

For the PSA, we extracted phylogenetic distances between species (e.g. time since species divergence) from the mega-tree GBOTB.extended.tre using the V.PhyloMaker package (Jin and Qian, 2019). We then performed phylogenetic signal analysis using the physig function from the phytools package (Revell, 2012), employing Pagel’s 𝜆 statistic (Pagel, 1999) as our measure of phylogenetic signal. This approach was appropriate for our dataset because Pagel’s 𝜆 can effectively handle polytomies —a convergent point (node) in a phylogeny where more than two lineages descend from a single ancestral lineage— which were observed among the Macleania species in our phylogenetic tree (Münkemüller et al., 2012).

#### 2.5.4 Crossed Mixed-Effect Models

Prior to model construction, we conducted Pearson correlations among all environmental variables (Figure S 2) using the cor.test function from the R stats package (R Core Team, 2013) to identify potential multicollinearity issues. This preliminary analysis was necessary for avoiding bias in model coefficients that could arise from highly correlated predictor variables (Schielzeth et al., 2020).

Based on correlation results, we constructed eight crossed mixed-effects models to test the effects of temperature (°C: mean/standard deviation/range), relative humidity (%: mean/standard deviation/range), and mean day length (hours) on spectrally predicted traits. All models incorporated crossed random effects for individuals and collection dates to account for repeated measurements and temporal variation, while environmental variables were included as fixed effects. The decision to include taxonomic level as an additional random factor was informed by phylogenetic signal analysis results described in Section 2.5.3.

Our modeling approach involved testing different aspects of environmental variability across the eight models. Models 1-5 examined the effects of mean temperature, mean relative humidity, mean day length, and their interactions on plant traits. Models 6-8 specifically focused on environmental variability, testing the effects of temperature standard deviation, relative humidity standard deviation, and the combined ranges of temperature and relative humidity, respectively (Table 2). All models were fitted using the lmer function from the lme4 package (Bates et al., 2015), ensuring appropriate handling of the nested random effects structure.

**Table 2.** The eight crossed mixed effect model equations used to test whether the environment had an effect over the spectrally derived traits which underlie spectra. Where 𝑦_𝑖𝑗𝑘_ is the spectrally derived trait, the β are the fixed-effects regression coefficients for the fixed effects T = Temperature (℃), RH = Relative Humidity (%), and DL = daylength (hours). The random complements to the β are 𝜇(𝑖) and 𝜈(𝑗) for the crossed random effects (Individuals and Year). Finally, 𝜀_𝑖𝑗𝑘_ is the unexplained variance of each model.

| MODEL | MODEL SPECIFICATION |
| --- | --- |
| M<sub>1</sub> | $y_{ijk} = \beta_0 + \beta_1 T_{mean} + u(i) + v(j) + \varepsilon_{ijk}$ |
| M<sub>2</sub> | $y_{ijk} = \beta_0 + \beta_1 T_{mean} + \beta_2 RH_{mean} + u(i) + v(j) + \varepsilon_{ijk}$ |
| M<sub>3</sub> | $y_{ijk} = \beta_0 + \beta_1 T_{mean} + \beta_2 RH_{mean} + \beta_3 DL_{mean} + u(i) + v(j) + \varepsilon_{ijk}$ |
| M<sub>4</sub> | $y_{ijk} = \beta_0 + \beta_1 T_{mean} + \beta_2 RH_{mean} + \beta_3 DL_{mean} + \beta_4 T_{mean} * RH_{mean} + u(i) + v(j) + \varepsilon_{ijk}$ |
| M<sub>5</sub> | $y_{ijk} = \beta_0 + \beta_1 T_{mean} + \beta_2 RH_{mean} + \beta_3 DL_{mean} + \beta_4 T_{mean} * RH_{mean} * DL_{mean} + u(i) + v(j) + \varepsilon_{ijk}$ |
| M<sub>6</sub> | $y_{ijk} = \beta_0 + \beta_1 T_{sd} + u(i) + v(j) + \varepsilon_{ijk}$ |
| M<sub>7</sub> | $y_{ijk} = \beta_0 + \beta_1 RH_{sd} + u(i) + v(j) + \varepsilon_{ijk}$ |
| M<sub>8</sub> | $y_{ijk} = \beta_0 + \beta_1 T_{range} + \beta_2 RH_{range} + u(i) + v(j) + \varepsilon_{ijk}$ |

When models exhibited non-normally distributed residuals explained by each random effect, via the Flinger-Killeen test, we implemented the BoxCoxME function from the tramME package to ensure compliance with model assumptions (Tamási and Hothorn, 2021). This modeling approach assumes a Standard Gaussian error distribution and applies a Bernstein basis transformation to the response variable (spectral trait data), which utilizes Bernstein polynomials to establish a flexible, non-parametric transformation function. This approach allows the transformation to adapt to the specific distributional characteristics of our data rather than constraining it to a predetermined parametric form.

## 3 Results

### 3.1 Classification of leaf spectra at different taxonomic levels considering temporal variation

Classification performance differed dramatically between the Stratified and Split models across all taxonomic levels (Table 3). The stratified model, trained on all three time periods (spring 2019, summer 2020, winter 2021), achieved high accuracy: genus-level classification performed best (97.5%, κ = 0.97), followed by family-level (93.1%, κ = 0.92) and species-level (88.4%, κ = 0.88). High kappa (κ) values indicate performance substantially better than random chance, with balanced precision, recall, and F1 scores across taxonomic ranks.

**Table 3.** Classification (Partial Least Squares-Discriminant Analysis) analysis results for the Split partition (model), where 2019 leaves were used as training and 2020 and 2021 and the Stratified partition (model) where all leaves were classified together and partitioned considering collection date, at the family (Family Split & Family Stratified), genus (Genus Split & Genus Stratified), and species (Species Split & Species Stratified) levels. All training models were cross validated using a bootstrap resampling to obtain the number of components.

| Model Type | Taxonomic Level | Number of Components | Accuracy | Kappa | Macro Precision | Macro Recall | Macro F1 | Weighted Precision | Weighted Recall | Weighted F1 |
| --- | --- | --- | --- | --- | --- | --- | --- | --- | --- | --- |
| SPLIT | Family | 30 | 0.35 | 0.21 | 0.62 | 0.35 | 0.42 | 0.35 | 0.78 | 0.41 |
| SPLIT | Genus | 19 | 0.37 | 0.22 | 0.37 | 0.29 | 0.52 | 0.37 | 1.00 | 0.53 |
| SPLIT | Species | 30 | 0.09 | 0.06 | 0.16 | 0.09 | 0.25 | 0.09 | 0.79 | 0.14 |
| STRATIFIED | Family | 30 | 0.93 | 0.92 | 0.94 | 0.94 | 0.94 | 0.93 | 0.93 | 0.93 |
| STRATIFIED | Genus | 22 | 0.98 | 0.97 | 0.98 | 0.97 | 0.97 | 0.98 | 0.98 | 0.98 |
| STRATIFIED | Species | 30 | 0.88 | 0.88 | 0.92 | 0.90 | 0.89 | 0.88 | 0.94 | 0.89 |

The split model, trained only on 2019 data and tested on 2020-2021 data, showed poor temporal generalization. Accuracy dropped to 34.7% (κ = 0.21) for family-level, 36.7% (κ = 0.22) for genus-level, and only 8.7% (κ = 0.06) for species-level classification.

Component analysis revealed taxonomic-specific complexity requirements. Family and species classifications used the maximum 30 components, indicating high spectral complexity needs without overfitting (confirmed by 1000 bootstrap iterations). Genus-level classification required fewer components (19-22), suggesting optimal complexity was reached within the tested range. The difference between stratified (22) and split (19) models reflects additional complexity needed for temporal variation.

The split model showed systematic misclassification bias. At the family level, 52% of non-Ericaceae samples were misclassified as Ericaceae (Figure 1A). At the genus level, 42% of samples were misclassified as Iochroma and 35% as Macleania (Figure 1C). At the species level, *M. coccoloboides* and *Osa pulchra* drove misclassifications, attracting 29.3% and 28.5% of samples, respectively (Figure 1E).

**Figure 1.**
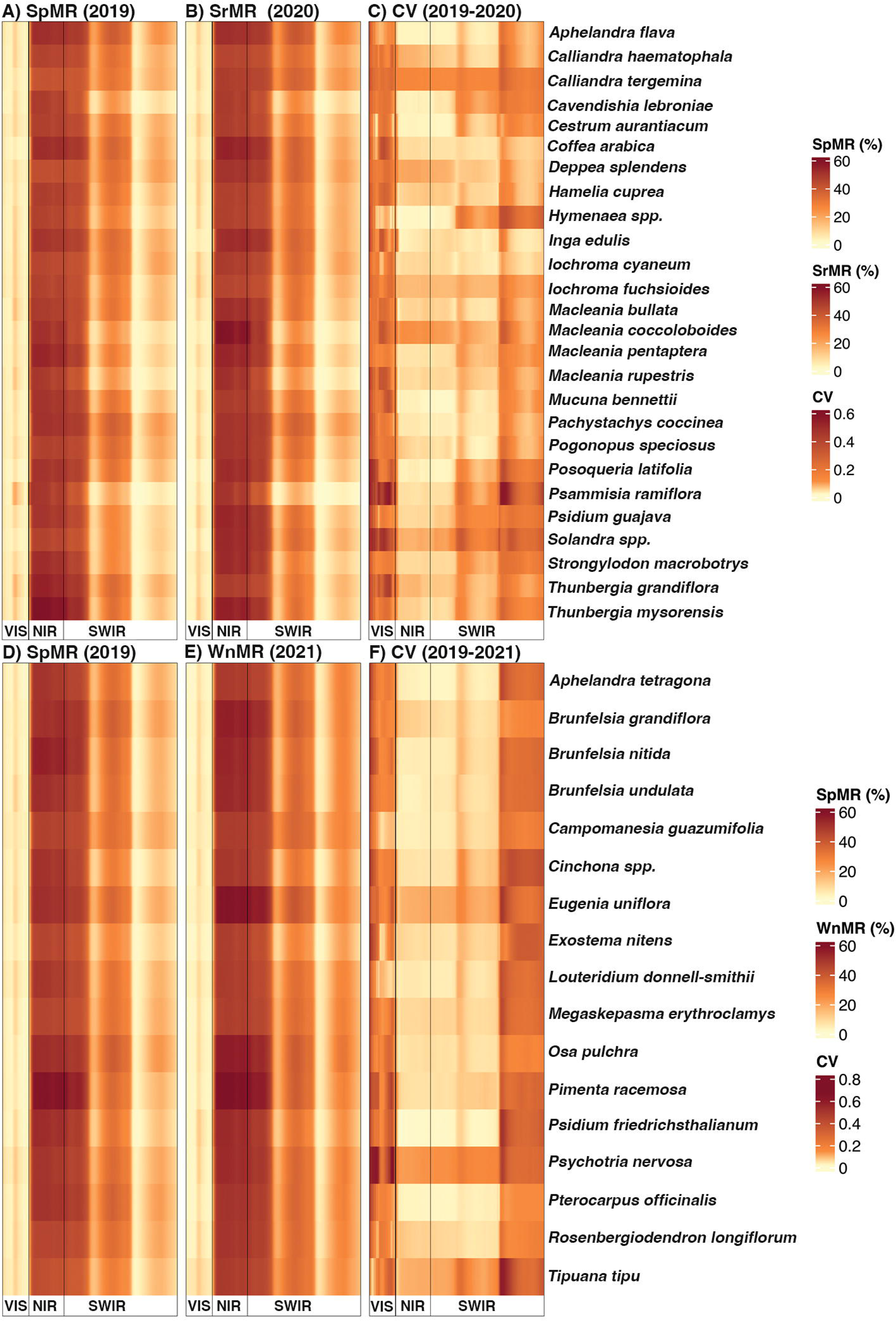
PLS-DA confusion matrices for Split and Stratified classification analyses at family (A, B), genus (C, D), and species (E, F) taxonomic levels. Split models (A, C, E) were trained using 2019 leaf samples and validated on independent 2020 and 2021 samples. Stratified models (B, D, F) were trained and tested using stratified random partitioning (70% training/30% testing).

The stratified model eliminated systematic bias. Family-level errors occurred only between taxonomically related families: Fabaceae-Myrtaceae (both Rosids) and Solanaceae-Rubiaceae (both Asterids). Genus-level errors were minimal, occurring only between Aphelandra and Thunbergia (both Acanthaceae) (Figure 1D). At the species level, only seven species showed misclassification, primarily among taxonomically related taxa. For example, 50% of *Brunfelsia undulata* samples were confused with *B. nitida*, and *Calliandra haematocephala* was distributed equally among itself (33%), *C. tergemina* (33%), and *Campomanesia guazumifolia* (33%) (Figure 1F).

### 3.2 Detection of variability within spectral reflectance

Seasonal spectral variability showed distinct patterns across wavelength regions (Figure 2). Coefficient of variation analysis between spring baseline measurements and summer/winter remeasurements revealed consistent trends across species, with the visible spectrum exhibiting the highest variability. In this region, winter-remeasured species displayed CV values ranging from 23.2% to 66.2% with the highest variation on the red region (653-675 nm) with an overall variation of 64.4%, while summer-remeasured species showed narrower but still substantial variation (CV = 29-62%), with the violet/blue region (400-405 nm) with an overall variation of 60.2%. Red edge variability differed between seasons: summer species peaked at 693 nm (CV = 51%), while winter species peaked at 680 nm with a maximum CV of 60%. In contrast, the near-infrared region (700-1100 nm) exhibited the lowest seasonal variability for both groups with an overall minimum variation of 1.2% for the winter-remeasured species and 1.5% for the summer-remeasured species.

**Figure 2.**
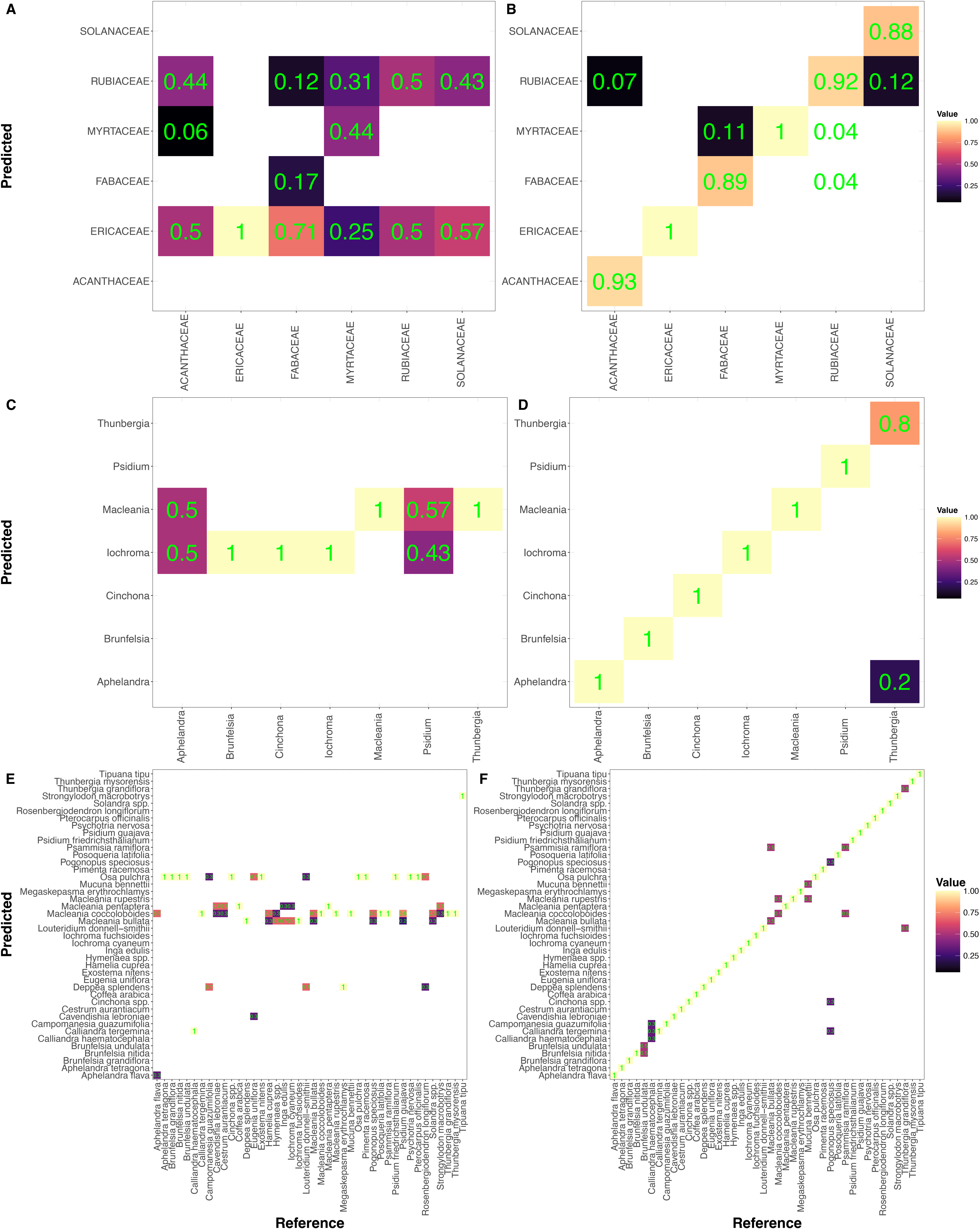
Mean spectral reflectance (MR%) across visible (VIS) to shortwave infrared (SWIR) wavelengths for baseline species sampled in spring 2019 (Sp; panels A & D, n=5 leaves per species) and species remeasured in summer 2020 (Sr; panel B, n=3 leaves per species) and winter 2021 (Wn; panel E, n=3 leaves per species). Coefficient of variation (CV%) calculated from baseline and remeasured leaves for 2020 (panel C) and 2021 (panel F). Spectral regions: VIS = 400-700 nm, NIR = 700-1100 nm, SWIR = 1100-2400 nm.

### 3.3 Prediction and temporal variation of plant chemical and structural traits

All leaf traits predicted for the NYBG leaf samples using PLSR coefficients obtained from the NEON foliar and spectroscopic datasets showed statistically significant change between the individuals remeasured in the winter of 2021(p-value_all_traits_ < 0.01), but not in the individuals remeasured in the summer of 2020 (p-value_all_traits_ > 0.05), with a much stronger effect on the individuals remeasured in the winter of 2020 (Table 4, Figure 3 C,D, Table S 2). Based on these results and the difference in the spectral behavior of this samples based on their identity, season, and year of remeasurement also observed in the LDA plot (Figure 3B) prompted us to run the linear mixed-effect models separately.

**Figure 3.**
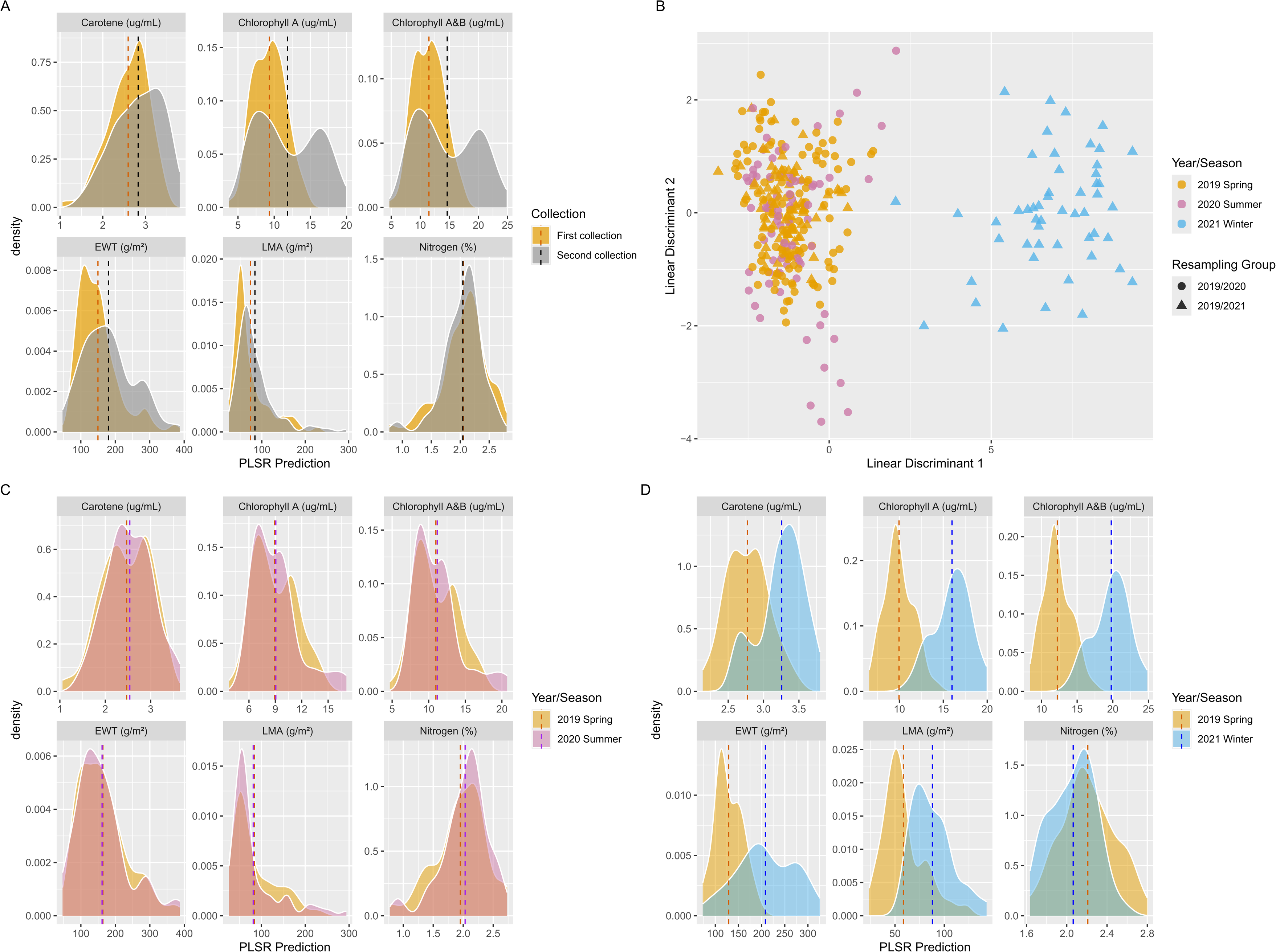
PLSR-predicted trait variation between seasons and linear discriminant analysis. Dashed lines indicate trait means. (A) Trait density distributions for baseline collection (2019) and second collections (2020 and 2021 combined). (B) First two linear discriminant variables showing similarities and differences between baseline (2019) and second collections (2020 and 2021). (C) Trait density distributions comparing baseline collection (2019) to summer 2020 samples. (D) Trait density distributions comparing baseline collection (2019) to winter 2021 samples.

**Table 4.** Results of Statistical Analysis of Temporal Trait Variation. Baseline measured conducted in the Spring of 2019.

| Resampling Group | Trait | N Pairs | Mean Difference | t-statistic | p-value | 95% CI Lower | 95% CI Upper | Effect Size (Cohen's d) |
| --- | --- | --- | --- | --- | --- | --- | --- | --- |
| Summer 2020 | Carotenoids ($\mu\text{g/mL}$) | 25 | 0.0871 | 1.821 | 0.09 | -0.01 | 0.19 | 0.36 |
| | Chl. A ($\mu\text{g/mL}$) | 25 | 0.2214 | 0.732 | 0.47 | -0.40 | 0.9 | 0.15 |
| | Chl. A&B ($\mu\text{g/mL}$) | 25 | 0.2856 | 0.779 | 0.44 | -0.47 | 1.04 | 0.16 |
| | EWT ($\text{g/m}^2$)* | 25 | -4.7364 | 152 | 0.79 | -13.02 | 7.17 | -0.19 |
| | LMA ($\text{g/m}^2$) | 25 | -3.8815 | -0.859 | 0.4 | -13.21 | 5.45 | -0.17 |
|  | Nitrogen (%) | 25 | 0.0911 | 2.805 | **0.01** | 0.02 | 0.16 | 0.56 |
| Winter 2021 | Carotenoids ($\mu\text{g/mL}$) | 17 | 0.4841 | 12.839 | **0** | 0.40 | 0.56 | 3.11 |
| | Chl. A ($\mu\text{g/mL}$)* | 17 | 6.037 | 153 | **0** | 5.41 | 6.56 | 6.23 |
| | Chl. A&B ($\mu\text{g/mL}$) | 17 | 7.5598 | 26.92 | **0** | 6.97 | 8.16 | 6.53 |
| | EWT ($\text{g/m}^2$) | 17 | 79.4006 | 7.574 | **0** | 57.18 | 101.62 | 1.84 |
| | LMA ($\text{g/m}^2$) | 17 | 29.4456 | 8.03 | **0** | 21.67 | 37.22 | 1.95 |
|  | Nitrogen (%) | 17 | -0.1454 | -4.813 | **2e-04** | -0.21 | -0.08 | -1.17 |

\* Indicates that the comparison between means was carried out with a Wilcoxon signed-rank test

### 3.4 Phylogenetic Signal Analysis

When we tested for the effect of the phylogeny over trait variation, measured as the interquartile range of the different estimated traits, we found inconclusive results for most traits, but EWT, since we could not reject the null hypothesis that the phylogenetic signal was different from zero for any of the two remeasuring events (p-value > 0.05; Table 5). EWT showed a statistically significant phylogenetic signal for the individuals remeasured in the Summer of 2020. Since these samples showed an influence of the evolutionary history of this trait in these species, we included the taxonomic rank family as a random effect on the crossed mixed-effect models to account for the lack of phylogenetic independence.

**Table 5.** Results for phylogenetic signal analysis of the interquartile range of all traits, predicted from the spectral reflectance, for both the species remeasured during the summer and winter using a trimmed angiosperm mega phylogeny and Pagel’s lambda. Baseline measured conducted in the Spring of 2019.

| Resampling Group | Trait | $\lambda$ | p-value |
| --- | --- | --- | --- |
| 2020 Summer | Chlorophyll A&B ($\mu g/mL$) | 7.49e-05 | 1 |
| | Chlorophyll A ($\mu g/mL$) | 7.49e-05 | 1 |
| | Carotenoids ($\mu g/mL$) | 7.49e-05 | 1 |
| | Equivalent Water Thickness ($g/m^2$) | 9.70e-01 | **2.51e-02** |
| | Leaf Dry Mass per Area ($g/m^2$) | 7.49e-05 | 1 |
|  | Nitrogen (%) | 7.49e-05 | 1 |
| 2021 Winter | Chlorophyll A&B ($\mu g/mL$) | 7.69e-05 | 1 |
| | Chlorophyll A ($\mu g/mL$) | 2.30e-01 | 8.93e-01 |
| | Carotenoids ($\mu g/mL$) | 2.83e-01 | 6.76e-01 |
| | Equivalent Water Thickness ($g/m^2$) | 5.06e-02 | 8.59e-01 |
| | Leaf Dry Mass per Area ($g/m^2$) | 2.12e-01 | 7.21e-01 |
|  | Nitrogen (%) | 7.69e-05 | 1 |

### 3.5 Influence of the environment over the optical properties of leaves

Summer 2020 remeasurements: Mixed-effects models were successful for all transformed traits except total chlorophyll and chlorophyll A, which violated model assumptions (Table 6). Average temperature and day length significantly affected carotenoid content (p < 0.01). EWT was influenced by average temperature, day length, average relative humidity, and the humidity × day length interaction (p < 0.05). LMA responded to the temperature × day length interaction (p < 0.01). Finally, Temperature range significantly affected leaf nitrogen content (p < 0.01).

**Table 6.** Results for most parsimonious crossed mixed effect models testing for the effect of the Conservatory environment over the spectrally BoxCox transformed predicted traits, for 25 species remeasured in the summer of 2020 based on AICc. The environmental variables presented in these models were T_max_ = Mean Temperature (℃), RH_mean_ = Mean Relative Humidity (%), DL_mean_ = Mean Daylength (hours), RH_range_ = Relative Humidity range (%), T_range_ = Temperature Range (℃), and RH_sd_ = Standard Deviation of the Relative Humidity (%). Baseline measured conducted in the Spring of 2019.

| Resampling Group | PLSR Trait | Environmental Variable | Estimate | Std. Error | z value | Pr(>\|z\|) |
| --- | --- | --- | --- | --- | --- | --- |
| Summer 2020 | Carotenoids ($\mu\text{g/mL}$) | $T_{\text{mean}}$ ($^{\circ}\text{C}$) | -2.144 | 0.588 | -3.644 | **2.68E-04** |
| | | $RH_{\text{mean}}$ (%) | -0.126 | 0.533 | -0.237 | 0.812 |
| | | $DL_{\text{mean}}$ (h) | 2.086 | 0.450 | 4.632 | **3.62e-06** |
| | | $T_{\text{mean}}$ ($^{\circ}\text{C}$) * $RH_{\text{mean}}$ (%) | -0.334 | 0.643 | -0.520 | 0.603 |
| | | $T_{\text{mean}}$ ($^{\circ}\text{C}$) * Mean Day Length (h) | 0.865 | 0.266 | 3.252 | **0.001** |
| | | $RH_{\text{mean}}$ (%) * Mean Day Length (h) | 0.483 | 0.327 | 1.474 | 0.141 |
| | | $T_{\text{mean}}$ ($^{\circ}\text{C}$) * $RH_{\text{mean}}$ (%) * $DL_{\text{mean}}$ (h) | 0.170 | 0.578 | 0.293 | 0.769 |
| | EWT ($\text{g/m}^2$)* | $T_{\text{mean}}$ ($^{\circ}\text{C}$) | 1.727 | 0.596 | 2.896 | **0.004** |
| | | $RH_{\text{mean}}$ (%) | -1.075 | 0.528 | -2.034 | **0.042** |
|  |  | Mean Day Length (h) | -1.224 | 0.444 | -2.756 | **0.006** |
| | | $T_{\text{mean}}$ ($^{\circ}\text{C}$) * $RH_{\text{mean}}$ (%) | -0.469 | 0.635 | -0.739 | 0.460 |
| | | $T_{\text{mean}}$ ($^{\circ}\text{C}$) * $DL_{\text{mean}}$ (h) | -1.624 | 0.276 | -5.877 | **4.19e-09** |
| | | $RH_{\text{mean}}$ (%) * $DL_{\text{mean}}$ (h) | -0.704 | 0.329 | -2.140 | **0.032** |
| | | $T_{\text{mean}}$ ($^{\circ}\text{C}$) * $RH_{\text{mean}}$ (%) * $DL_{\text{mean}}$ (h) | -0.386 | 0.570 | -0.678 | 0.498 |
| | LMA ($\text{g/m}^2$) | $T_{\text{mean}}$ ($^{\circ}\text{C}$) | -0.099 | 0.576 | -0.171 | 0.864 |
| | | $RH_{\text{mean}}$ (%) | -0.408 | 0.529 | -0.772 | 0.440 |
| | | $DL_{\text{mean}}$ (h) | 0.144 | 0.436 | 0.330 | 0.742 |
| | | $T_{\text{mean}}$ ($^{\circ}\text{C}$) * $RH_{\text{mean}}$ (%) | 0.335 | 0.635 | 0.528 | 0.598 |
| | | $T_{\text{mean}}$ ($^{\circ}\text{C}$) * $DL_{\text{mean}}$ (h) | -0.768 | 0.258 | -2.980 | **0.003** |
| | | $RH_{\text{mean}}$ (%) * $DL_{\text{mean}}$ (h) | -0.425 | 0.327 | -1.298 | 0.194 |
| | | $T_{\text{mean}}$ ($^{\circ}\text{C}$) * $RH_{\text{mean}}$ (%) * $DL_{\text{mean}}$ (h) | -0.370 | 0.574 | -0.645 | 0.519 |
| | Nitrogen (%) | $RH_{\text{range}}$ (%) | -0.032 | 0.137 | -0.236 | 0.813 |
| | | $T_{\text{range}}$ ($^{\circ}\text{C}$) | 0.334 | 0.086 | 3.898 | **9.69e-05** |

\*EWT model included Family as a random effect.

Winter 2021 remeasurements: All transformed traits, except EWT, met model assumptions (Table 7). All photosynthetic pigments (chlorophyll A, total chlorophylls, carotenoids) were significantly affected by average temperature, relative humidity, and day length (p < 0.05). LMA responded to day length and its interactions with temperature and relative humidity (p ≤ 0.05). Leaf nitrogen was influenced by the relative humidity × day length interaction (p < 0.05).

**Table 7.**
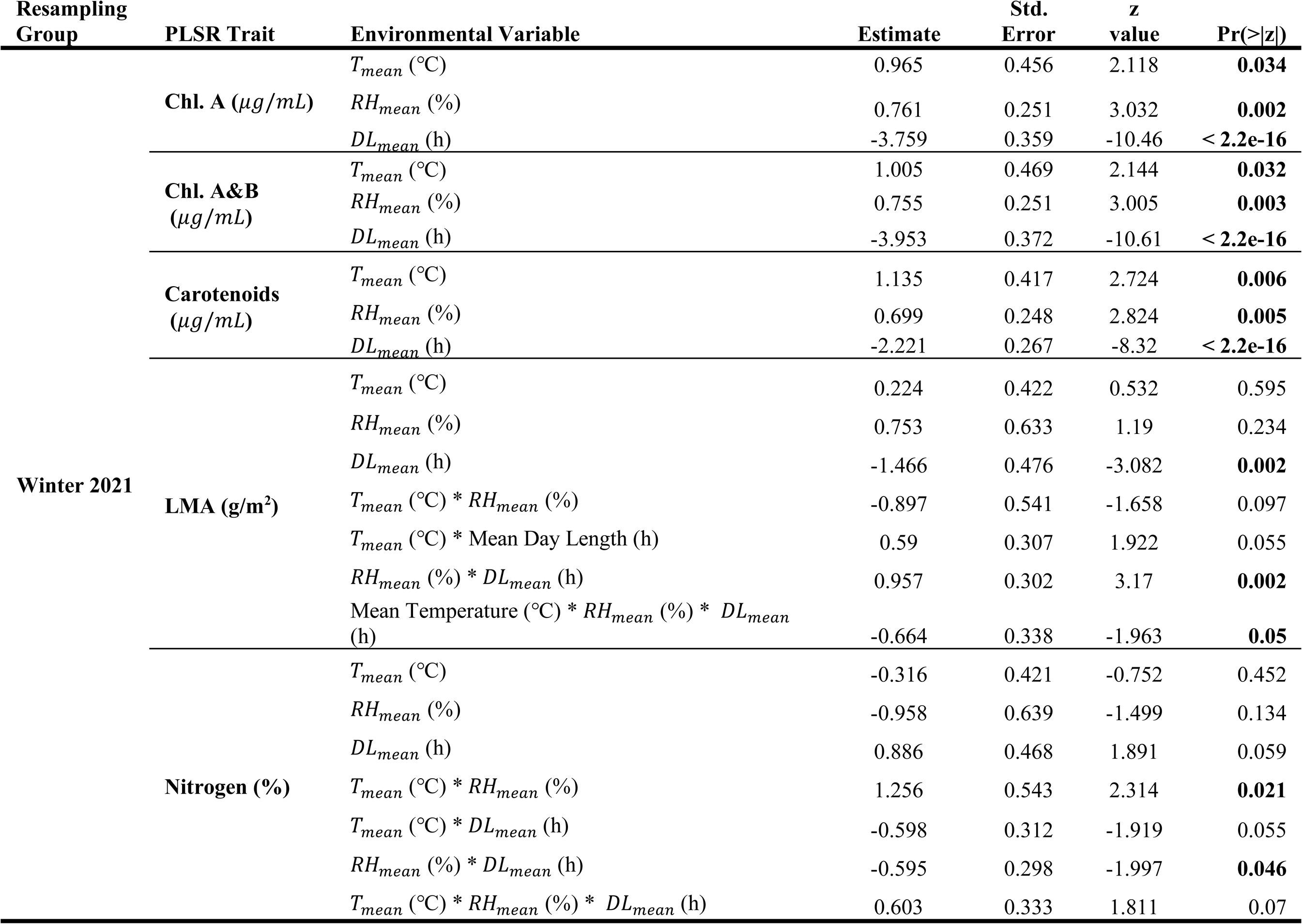
Results for most parsimonious mixed effect models testing for the effect of the Conservatory environment over the spectrally predicted traits, for 16 species remeasured in 2021 based on AIC. All traits were BoxCox transformed. The environmental variables presented in these models were Tmax = Mean Temperature (℃), RHmean = Mean Relative Humidity (%), and DLmean = Mean Daylength (hours). Baseline measured conducted in the Spring of 2019.

### 3.6 Use of Artificial Intelligence (AI)

We used Goddard Space Flight Center’s in-house AI tool (ChatGSFC) to optimize PLSR coefficient extraction from the NEON datasets (Quinteros Casaverde and Serbin, 2025), to troubleshoot code, and to improve the clarity of the manuscript.

## 4 Discussion

Our results demonstrate that temporal spectral variability significantly impacts the reliability of leaf-based taxonomic classification. The stratified model, incorporating multi-seasonal data, achieved high accuracy across all taxonomic levels (κ > 0.9) compared to temporally limited split models (κ < 0.5). This finding is particularly relevant given the increasing availability of continuous spectral data from orbital sensors like NASA’s EMIT (Green et al., 2020; Thompson et al., 2024)and PACE (Gorman et al., 2019) , which capture year-round phenological variation rather than single-season snapshots.

While most leaf-level spectral classification studies rely on single-season collections, recent airborne and spaceborne research has begun addressing temporal variability challenges. Studies using aerial and satellite imaging spectroscopy have successfully integrated temporal spectral variation for species classification using advanced statistical methods(Jiang et al., 2025; Xu et al., 2021; Zhao et al., 2023). Our leaf-level results confirm that methods capable of handling spectral collinearity, such as Partial Least Squares Discriminant Analysis, can effectively integrate temporal variability across taxonomic levels, supporting the scalability of these approaches from laboratory to landscape applications.

A closer assessment into the spectral variability patterns revealed underlying physiological drivers of temporal variation. The prominent seasonal changes in the violet/blue region (400-405 nm; CV_summer20_ up to 60%), the red region (653-675 nm; CV_winter21_ up to 64%), and the variable red edge positions (summer: CV_693nm_ = 51% and winter: CV_680nm_ = 60%) reflect dynamic photosynthetic pigment responses to environmental conditions (Figure 2 and Figure 3), consistent with previous tropical (Hesketh & Sánchez-Azofeifa, 2012) and temperate (Schweiger et al., 2018) studies. Conversely, the stability of near-infrared regions (700 - 1000 nm) indicates that structural properties remain more consistent than biochemical components across seasons (Gates et al., 1965; Slaton et al., 2001).

The misclassification patterns in the stratified model revealed phylogenetic constraints on spectral distinctiveness (Figure 3 B,D,E). Errors occurred primarily between taxonomically related taxa (e.g., Solanaceae-Rubiaceae, both Asterids), suggesting that evolutionary relationships impose limits on spectral discrimination. This aligns with documented phylogenetic signals in plant spectral properties, particularly within Asterids (Meireles et al., 2020).

The results also showed that environmental factors drove trait-specific responses that varied seasonally. Summer measurements showed consistent temperature and humidity effects across transformed predicted traits, while winter measurements invariably included day length effects (Table 6 and Table 7). The observed relationships between environmental variables and transformed predicted structural and biochemical leaf traits demonstrated that spectral variation also reflects adaptive and non-linear physiological responses to fluctuating conditions within the conservatory environment as observed in nature (Huxley et al., 2023; Niklas et al., 2007; Poorter et al., 2010), contributing to misclassification.

Conducting this research within the controlled environment of the Enid A. Haupt Conservatory (EAHC) at NYBG provided both opportunities and limitations for understanding taxonomic misclassification causes. While the EAHC’s semi-controlled environment represents a simplified ecological system lacking environmental stochasticity, biotic interactions, and resource gradients that drive plant stress in natural settings, it enabled us to: (1) isolate seasonal environmental effects (temperature, humidity, daylength) without confounding factors, (2) maintain consistent sampling protocols, (3) ensure taxonomic identity certainty, and (4) conduct replicated measurements under standardized conditions.

We acknowledge that field-based spectral studies offer complementary advantages our controlled approach cannot provide, particularly the spatial and temporal scales relevant to satellite and airborne remote sensing applications. However, the controlled conservatory environment provided essential baseline understanding of how environmental variation affects plant spectral signatures (Serbin et al., 2019), establishing the mechanistic foundation for environmental effects on spectral taxonomy. These findings can guide comprehensive field validation studies and inform robust spectral classification methods for operational remote sensing applications.

## 5 Conclusions

In conclusion, this study establishes that temporal representativeness is fundamental for reliable spectral-based plant taxonomy. The dramatic performance differences between temporally comprehensive versus limited training datasets highlight the critical importance of capturing seasonal variation in spectral signatures. Genus-level classification emerged as the optimal balance between taxonomic resolution and temporal stability, achieving 97.5% accuracy while requiring moderate model complexity.

The integration of environmental data revealed that spectral variability is not random noise but reflects meaningful physiological responses to changing conditions. This understanding provides a pathway for improving spectral classification by incorporating environmental context into models. For operational applications using continuous satellite monitoring, our findings suggest that successful plant identification systems must account for phenological cycles and environmental variation rather than relying on single-season spectral libraries.

Future research should focus on developing adaptive classification frameworks that can leverage temporal spectral patterns as informative features rather than treating them as obstacles to overcome. This approach could transform spectral remote sensing from a static taxonomic tool into a dynamic system capable of monitoring both species identity and physiological status across landscapes and seasons.

## Acknowledgments

We thank Drs. A. Berkov, A.C. Carnaval, K. Dexter, and F.A. Michelangeli for manuscript revisions, and Drs. A. Schiklomanov and D. Kathuria for revision feedback. We appreciate M. Hachadourian, C. Primeau, and the NYBG Enid A. Haupt Conservatory staff for accommodating our research during the COVID-19 pandemic. Thanks to L. Adhikary and K. Martinez (LaGuardia Community College Students) for field assistance, CUNY EEB graduate students for constructive feedback, and the CUNY Graduate Center and Southeastern Universities Research Association (award #80NSSC22M0001) for financial support.

## Notes

https://doi.org/10.5281/zenodo.18097029

### Competing Interest Statement

The authors have declared no competing interest.

### Summary of Updates

We implemented comprehensive revisions to strengthen the manuscript's scientific rigor and clarity. The title was revised to better reflect our key finding regarding environmental impacts on spectral taxonomy. The introduction was refined for improved clarity and conciseness. Substantial methodological improvements were made, including optimizing the Partial Least Squares Discriminant Analysis (PLSDA) by switching from pls to kernel-pls algorithm for enhanced robustness with small sample sizes, and adding additional model performance measures to the PLSDA results. We streamlined the trait prediction analysis by utilizing NEON leaf-level spectra and foliar traits to derive PLSR coefficients for all traits, accompanied by a newly published GitHub repository to ensure reproducibility. Statistical analyses were strengthened through the addition of linear discriminant analysis to demonstrate trait seasonality and both parametric and non-parametric pairwise t-tests to statistically validate seasonal changes in trait composition. A dedicated section addressing the utilization of artificial intelligence was incorporated. The results section was updated to include all new findings, while the discussion was restructured by separating conclusions from the main discussion for improved organization. Finally, the manuscript was enhanced with updated references and acknowledgements.

