## Supplementary figures and images for "Environmentally driven changes in leaf spectral signatures impact plant spectral taxonomy"

### Supplemental Figure 1

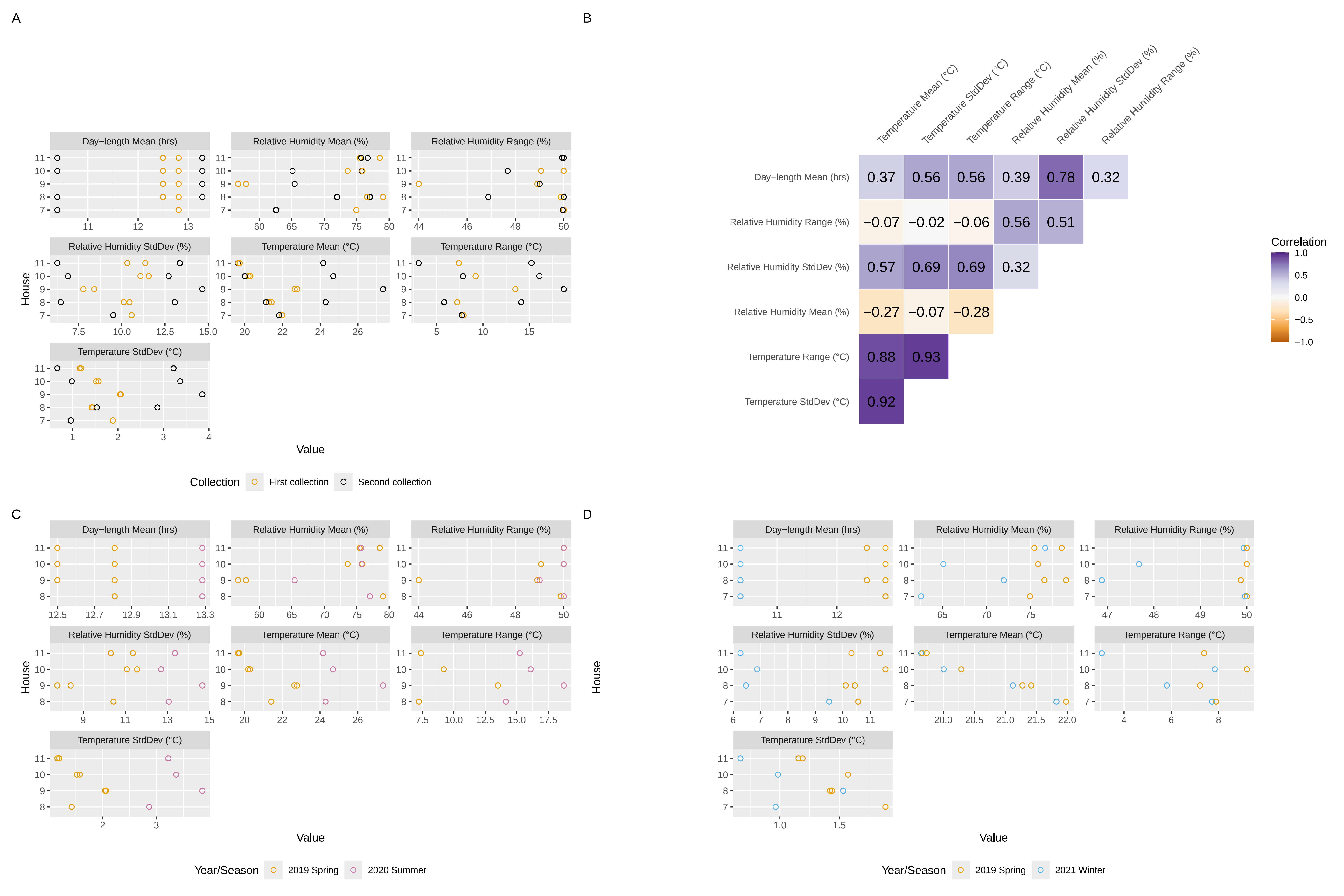

### Supplemental Figure 2

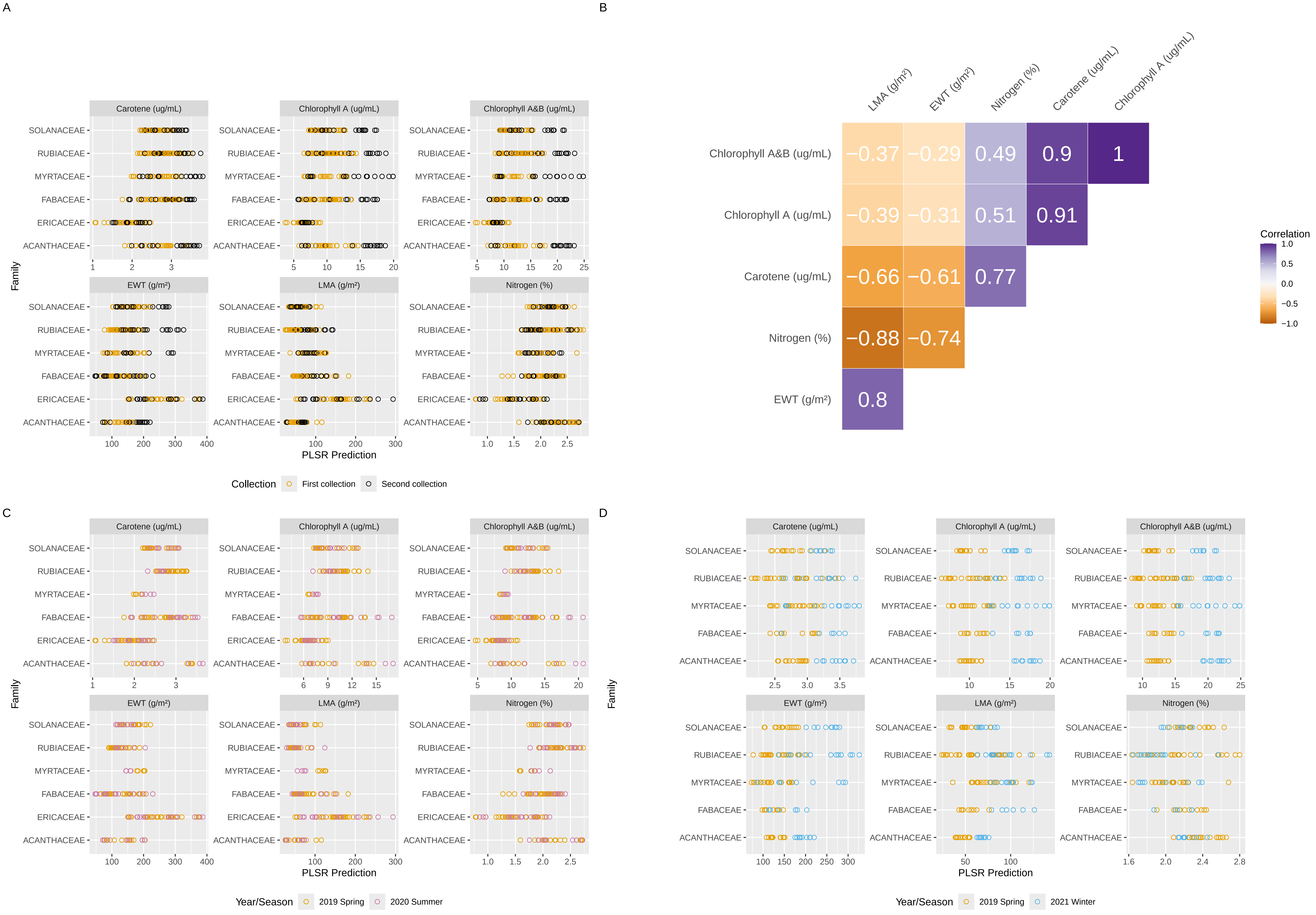
