## Supplemental Tables for "Environmentally driven changes in leaf spectral signatures impact plant spectral taxonomy"

SUPPLEMENTARY INFORMATION

Supplementary Tables

Table S 1. Species list. First column shows individual species collected, grouped by family. Column "2019" represents baseline collection. Columns "2020" and "2021" indicate whether species were remeasured in those years. Last column indicates family name.

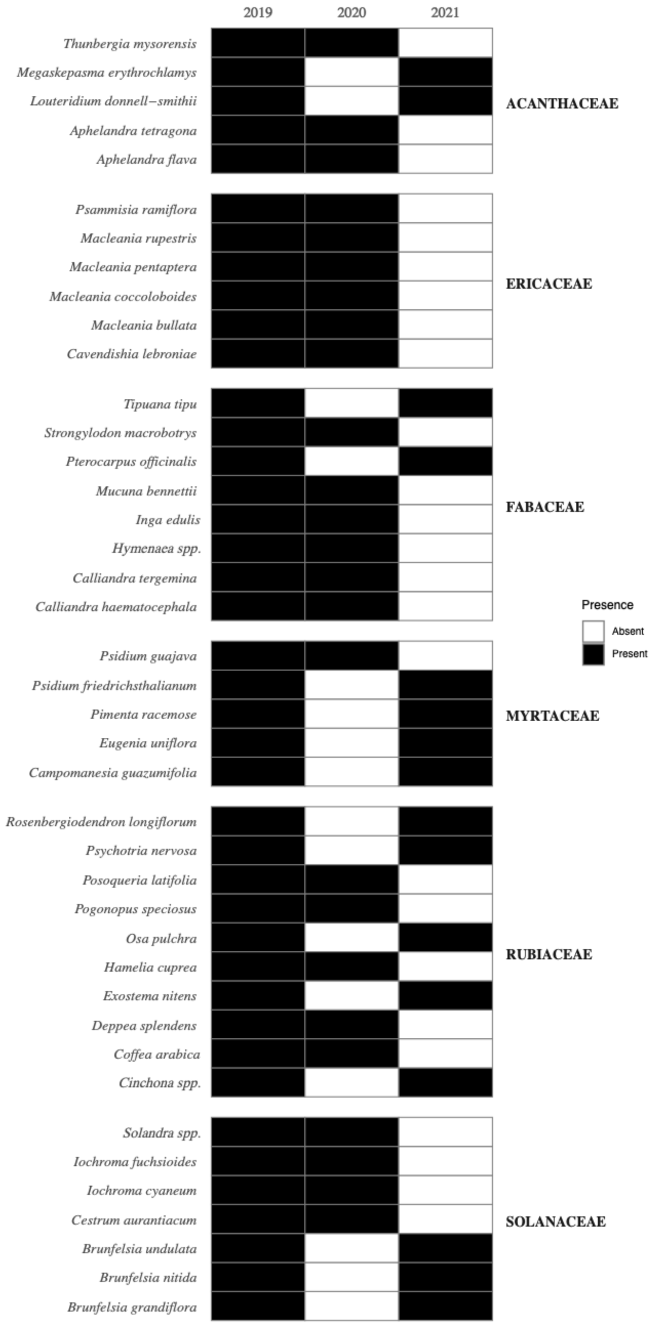

Table S 2. Model Performance Metrics for PLSDA.

| Metric | Range | Definition | Formula | Source |
|---|---|---|---|---|
| Cohen's Kappa | - 1 to 1 | Measures the agreement between the classifier and the observed classes, obtained by using the test dataset | $\frac{\mathrm{Pr(actual)} - \mathrm{Pr(expected)}}{1 - \mathrm{Pr(expected)}}$ | De Diego et al. 2022; Lantz, 2019 |
| Precision | 0 to 1 | Measures how often the model correctly predicting a true positive. | $\frac{N_{\text{true positives}}}{N_{\text{true positives}} + N_{\text{false positives}}}$ | Lantz, 2019 |
| Recall | 0 to 1 | Measures result completeness. | $\frac{N_{\text{true positives}}}{N_{\text{true positives}} + N_{\text{false negatives}}}$ | Lantz, 2019 |
| F1 score | 0 to 1 | Measure model performance as the harmonic mean between precision and recall. | $\frac{\text{2 x precision x recall}}{\text{precision} + \text{recall}}$ | Lantz, 2019 |

Table S 3 Thirty-day summary metrics for temperature (°C), relative humidity (%), and daylength (h) from the Enid A. Haupt Conservatory houses encompassing the Rain Forest Pavilion, used in the crossed mixed-effects models.

| Year | House | $T_{mean}$ | $T_{sd}$ | $T_{range}$ | $RH_{mean}$ | $RH_{sd}$ | $Rh_{range}$ | $DL_{mean}$ |
|---|---|---|---|---|---|---|---|---|
| 2019 | 7 | 21.89 | 1.91 | 7.95 | 71.69 | 10.58 | 50 | 12.5 |
| | 8 | 21.28 | 1.44 | 7.22 | 76.6 | 10.1 | 49.87 | 12.5 |
| | 9 | 22.64 | 2.04 | 13.5 | 56.73 | 7.77 | 48.9 | 12.5 |
| | 10 | 20.2 | 1.52 | 9.2 | 73.6 | 11.07 | 49.06 | 12.5 |
| | 11 | 19.66 | 1.16 | 7.39 | 75.45 | 10.31 | 50 | 12.5 |
| 2020 | 7 | 24.74 | 3.15 | 15.76 | 75.63 | 13.72 | 50 | 13.28 |
| | 8 | 24.29 | 2.87 | 14.13 | 77.04 | 13.06 | 50 | 13.28 |
| | 9 | 27.34 | 3.86 | 18.73 | 65.44 | 14.66 | 49 | 13.28 |
| | 10 | 24.69 | 3.37 | 16.1 | 75.78 | 12.71 | 50 | 13.28 |
| | 11 | 24.16 | 3.22 | 15.24 | 75.68 | 13.36 | 50 | 13.28 |
| 2021 | 7 | 21.83 | 0.96 | 7.72 | 62.58 | 9.51 | 49.96 | 10.39 |
| | 8 | 21.12 | 1.53 | 5.81 | 71.97 | 6.46 | 46.88 | 10.39 |
| | 9 | 21.17 | 2.21 | 9.84 | 67.97 | 6.1 | 43.99 | 10.39 |
| | 10 | 20.01 | 0.98 | 7.84 | 65.11 | 6.88 | 47.68 | 10.39 |
| | 11 | 19.64 | 0.67 | 3.06 | 76.68 | 6.26 | 49.93 | 10.39 |

### Supplementary Figures

Figure S 1. Environmental variable variation and correlation analysis between variables. (A) Differences between baseline collection (2019) and second collections (2020 and 2021). (B) Correlation analysis between environmental variables. (C) Variation between spring 2019 and summer 2020. (D) Variation between spring 2019 and winter 2021.

Figure S 2. PLSR trait variation and correlation analysis. (A) Variation of leaf traits across seasons and years for total chlorophylls (a&b), chlorophyll a, carotene density, equivalent water thickness (EWT), leaf dry mass per area (LMA), and nitrogen percent per family compared to baseline collection (2019). (B) Correlation analysis between PLSR traits. (C) Orange dots correspond to species measured in baseline collection (spring 2019), and red dots correspond to species remeasured in summer 2020. (D) Blue dots correspond to species remeasured in winter 2021.

Figure S 3. Mean spectral reflectance (fraction) from 400 nm to 2400 nm for six botanical families: Acanthaceae (ACA), Fabaceae (FAB), Ericaceae (ERI), Myrtaceae (MYR), Rubiaceae (RUB), and Solanaceae (SOLA). Spectral regions denoted are visible (VIS), near infrared (NIR), and shortwave infrared (SWIR).
